# Acetylcholine signaling in the medial prefrontal cortex mediates the ability to learn an active avoidance response following learned helplessness training

**DOI:** 10.1101/2023.09.23.559126

**Authors:** Zuhair I. Abdulla, Yann S. Mineur, Richard B. Crouse, Ian M. Etherington, Hanna Yousuf, Jessica J. Na, Marina R. Picciotto

## Abstract

Increased brain levels of acetylcholine (ACh) are observed in subsets of patients with depression and increasing ACh levels chronically can precipitate stress-related behaviors in humans and animals. Conversely, optimal ACh levels are required for cognition and memory. We hypothesize that ACh signaling is important for encoding both appetitive and stress-relevant memories, but that excessive increases in ACh result in a negative encoding bias in which memory formation of a stressful event is aberrantly strengthened, potentially contributing to the excessive focus on negative experience that could lead to depressive symptoms. The medial prefrontal cortex (mPFC) is critical to control the limbic system to filter exteroceptive cues and stress-related circuits. We therefore evaluated the role of ACh signaling in the mPFC in a learned helplessness task in which mice were exposed to repeated inescapable stressors followed by an active avoidance task. Using fiber photometry with a genetically-encoded ACh sensor, we found that ACh levels in the mPFC during exposure to inescapable stressors were positively correlated with later escape deficits in an active avoidance test in males, but not females. Consistent with these measurements, we found that both pharmacologically- and chemogenetically-induced increases in mPFC ACh levels resulted in escape deficits in both male and female mice, whereas chemogenetic inhibition of ACh neurons projecting to the mPFC improved escape performance in males, but impaired escape performance in females. These results highlight the adaptive role of ACh release in stress response, but also support the idea that sustained elevated ACh levels contribute to maladaptive behaviors. Furthermore, mPFC ACh signaling may contribute to depressive symptomology differentially in males and females.

## Introduction

Approximately ∼7.1% of the adult US population (17 million U.S. adults) experience at least one episode of major depression each year (NIMH 2017). Most currently available treatments for major depressive disorder (MDD) work by modulating monoaminergic and glutamatergic neurotransmitter systems, however these medications are effective for only two-thirds of patients (*1*) likely because depression is a multi-faceted disorders with differential etiology. It is therefore necessary to explore alternative mechanisms that might be involved in MDD or its treatment, such as the cholinergic system.

The hypercholinergic hypothesis of depression, first proposed by Janowsky and colleagues in 1972 (*2*), posited that elevated levels of acetylcholine (ACh) are responsible for depressive symptoms. Increased brain ACh levels are observed in those with active depression or a history of the disorder (*3–5*) and manipulating ACh transmission significantly alters stress-related behaviors. For instance, blocking breakdown of ACh with an acetylcholinesterase (AChE) antagonist in humans increases depressive symptoms and can reverse mania (*5, 6*). Similarly, AChE inhibition increases passive or maladaptive coping strategies in animals (*7–13*).

ACh signaling is also involved in cognition and reward-related behaviors (*14–16*), with ACh release in the prefrontal cortex (PFC) particularly critical for focused attention (*17*), indicating that optimal homeostatic levels of ACh are required for proper modulation of attention, learning, and mood (*18*). In response to the seemingly contrasting findings that ACh signaling is involved in both maladaptive responses to stress and adaptive cognitive processes, we hypothesize that in response to prolonged or inescapable stress, excessive increases in ACh signaling results in a negative encoding bias, thereby facilitating memory formation of the stressful event and/or improving recall and consolidation of these negative memories (*10*). This bias could contribute to the etiology of depression or result in depressive symptomology.

Here we evaluate the contribution of ACh signaling in the medial PFC (mPFC) to escape performance in a learned helplessness (LH) paradigm, in which male and female mice are repeatedly exposed to inescapable stressors and then tested in an active avoidance task, to evaluate coping behavior in response to stress. We used fiber photometry to measure ACh levels in mPFC during development of LH and determined whether these levels could be related to escape latency in the active avoidance task. We then evaluated the causal mechanisms underlying this behavior by manipulating ACh levels pharmacologically with the AChE inhibitor physostigmine, or chemogenetically to alter mPFC ACh release from cholinergic neurons projecting to the mPFC. These studies demonstrate that increasing ACh signaling in mPFC is sufficient to promote passive coping and induce escape deficits, supporting the possibility that ACh could contribute to the negative encoding bias observed in individuals with MDD.

## Methods

### Animals

Male and female, C57BL/6j mice (WT) were obtained from Jackson Labs (Bar Harbor, ME) at 8 weeks of age. ChAT-IRES-Cre (Δneo, ChAT-Cre; B6.129S-*Chat^tm1(cre)Lowl^*/MwarJ; stock/strain #031661) expressing male and female mice were bred in our facility for more than 10 generations and backcrossed onto the C57BL/6J background. Lines were maintained as heterozygotes (HET) and mice for this experiment were derived from HET x WT pairings. All mice were housed in group of 2 to 5 in microisolators equipped with an automatic watering system. Lights were on a 12:12 light:dark cycle, room temperature was maintained at 22.1 ± 1° C. Sterilized food and water were available *ad libitum*. All procedures were approved by the Yale Institutional Animal Care and Use Committee.

### Drugs

Physostigmine was purchased from Sigma-Aldrich. Clozapine-N-oxide dihydrochloride (CNO) was purchased from Hello Bio ((*19*); Princeton, NJ). For i.p. injections, physostigmine was dissolved in phosphate-buffered saline (PBS) at 0.015 mg/ml and delivered at 10 ml/kg bodyweight i.p for a dose of 0.15 mg/kg. CNO was dissolved in PBS at 0.7 mg/ml and delivered at 10 ml/kg bodyweight i.p for a dose of 7 mg/kg.

### Surgery

Surgeries for fiber photometry were carried out following established procedures (*14*). Briefly, 0.4 µl of a viral vector carrying a fluorescent ACh sensor (AAV9 hSyn-ACh4.3 (GACh3.0; WZ Biosciences Inc. (*20*)) was injected into the left hemisphere of the mPFC (Virus: AP +1.9, ML +0.4, DV −3.0) of C57BL/6J mice using a stereotaxic frame mounted with a 2 µl Hamilton Neuros syringe (Reno, NV, USA) at a rate of 0.1 µl/min. The needle was left in place for 5 min before the start of and after the conclusion of the injection. A fiber-optic cannula (Doric Lenses, Quebec City, Quebec, Canada) was implanted just above the injection site (Fiber: AP +1.9, ML +0.4, DV −2.95) and affixed to the skull using opaque dental cement (Parkell Inc., Edgewood, NY). Following surgery mice were allowed to recover for a minimum of 3 weeks to allow for adequate expression of GACh3.0.

Surgeries for chemogenetic experiments followed the same local infusion protocol as above except that 0.8 µl per hemisphere of a virus carrying a construct encoding a designer receptor exclusively activated by a designer drug (DREADD) or a control construct was injected bilaterally into the mPFC of ChAT Cre mice (AP +1.94, ML ±0.4, DV −3.1) using the following retrograde, Cre-dependent vector: pAAV-hSyn-DIO-HM3D(Gq)-mCherry (excitatory DREADD construct), pAAV-hSyn-DIO-HM3D(Gi)-mCherry (inhibitory DREADD construct), and pAAV-hSyn-DIO-mCherry (control construct; Addgene). Mice were allowed to recover for a minimum of 4 weeks after surgery to allow for adequate DREADD expression.

### Behavior

Learned helplessness (LH) methods were adapted from published studies (*21, 22*). LH is a three-day procedure occurring in two-chamber shuttle boxes with an automated gate between the chambers (MedAssociates, VT). This entire assembly was placed inside a sound attenuating chamber modified to allow for fiber photometry recordings. Mice received 120, 4-s inescapable shocks (0.3 mA) delivered semi-randomly (∼26 s inter-trial interval: ITI) over the course of 1 h during each of 2 induction trials, approximately 24 h apart. Control mice were placed in shock chambers for 1 h each day, but no shocks were administered. Roughly 24 h following induction trial 2, mice underwent active avoidance testing consisting of 30 trials of escapable shocks (0.3 mA) that terminated either upon escape or after 24 s, with an ITI of 10 s. A *k-means* clustering algorithm was used to define mice as “helpless” or “resilient” based on escape latency and escape failures, allowing for unbiased assignment of mice into two groups of maximal distinction. For DREADD experiments, mice were injected with 7 mg/kg CNO 30 min prior to each of the two induction trials, but not prior to active avoidance testing. For pharmacological experiments, 0.15 mg/kg physostigmine was administered 30 min prior to induction on day 2, but not prior to active avoidance testing.

### Fiber Photometry

Fiber photometry was used to measure ACh levels in the mPFC of behaving mice following established procedures (*14*). Briefly, fluorescent measurements of ACh were recorded using two standard Doric Lenses 1-site Fiber Photometry Systems, capable of detecting fluorescence at 405 nm (= “isosbestic”) and 465 nm (“signal”). A TTL adapter (Med Associates Inc.) was connected to each system to timestamp shock instances. Photometric signals were recorded using Doric Neuroscience Studio. Data were analyzed via custom written MATLAB code. First, baseline fluorescence (F0) was calculated using a first order least means squares regression over the course of the ∼1 h induction session. We then calculated the change in the fluorescence (ΔF) for any given timepoint as the difference between it and F0, divided by F0 and multiplied by 100 to obtain the %ΔF/F0 for the signal. Finally, the %ΔF/F0 was z-scored. Graphs show the averaged z-scored ΔF/F0 traces aligned to shock onset and the area under the curve (AUC) was derived from these averages.

### Tissue processing and verification of construct expression and targeting

Following completion of active avoidance testing, mice for fiber photometry and DREADD experiments were anesthetized with 0.2-0.3 mL Fatal Plus® (Patterson Veterinary Supplies, Inc., Devans, MA, USA) and perfused intracardially with 4% PFA for 2 min. Brains were removed and stored overnight at 4° C in 4% PFA for post-fixation and then in a 30% Sucrose in PBS solution for cryoprotection until they sank. Brains were then sliced at 40 µm thickness with a Leica SM2000R sliding microtome (Leica Biosystems, Wetzlar, Germany) and mounted on slides (Superfrost Plus Microscope Slides, Fisherbrand, Thermo Fisher Scientific, Waltham, MA, USA) and examined through an Olympus Fluoview FV10i confocal microscope. In fiber photometry experiments, GACh3.0 expression was confirmed using EGFP fluorescence in the mPFC and fiber tract was imaged to determine placement of the fiber. In DREADD experiments, mCherry fluorescence was used to confirm infection of the viruses in the mPFC and the nucleus basalis.

### Statistics

No outliers were removed from analyses; however, animals with off-target viral infusion or fiber misplacement were not included in analyses. On rare occasions, mice would cling to the walls of the shock chambers during active avoidance testing and their data were removed from all analyses. Statistical analyses were performed in GraphPad Prism 9 software. Values are expressed as mean ± standard error of the mean (SEM). Analysis of variance (ANOVA) was used when comparing multiple groups with an alpha of 95%. When ANOVA reached significance, *post hoc* multiple-comparisons were performed with Tukey’s range tests. Two-tailed Student’s t-tests were used when only two groups were compared. Two-sided Chi-square analyses were used for comparing helpless to resilient animals. Mann-Whitney U tests were used to compare number of escape failures between groups. Custom written MATLAB code was used for fiber photometry and *k-means* clustering in this study.

### Code

All code used in this study is freely available both online and upon request.

## Results

### Exposure to inescapable foot shocks produces subsequent escape deficits in an active avoidance task

Exposure to inescapable stressors reliably induces a learned helplessness phenotype in groups of mice, however individual animals can be either resilient (escaped quickly and often) or helpless (escaped slowly and seldom) to the stressor (*22–24*). In the current experiments we used a *k*-means clustering algorithm to define mice as “resilient” or “helpless” in an active avoidance task following two sessions of exposure to inescapable foot shock (*23*). Exposure to inescapable foot shock significantly increased escape latencies in the active avoidance test (Supplementary Fig. 1A; t (29) = 6.68, p < 0.0001) and the *k-means* clustering algorithm identified 2/3 of the mice in the shock group as helpless.

### ACh levels in the mPFC are predictive of escape behavior in learned helplessness

We next measured ACh in the mPFC of mice undergoing learned helplessness training (Fig. 1A) to determine whether ACh levels during training could predict behavior in later active avoidance testing. Given the invasive nature of stereotaxic surgery it was important to establish that viral infusion and fiber implantation did not disrupt learned helplessness behavior. Mice in the shock group had significantly increased escape latencies relative to naïve controls (Supplementary Fig. 1B; mean diff = 10.2 s, F (1,24) = 11.49, p < 0.01). Shock increased escape latencies in both male and female mice (Fig 1C; t (13) = 2.29, p<0.05 and t (11) = 2.48, p<0.05 respectively). Approximately the same number of male and female mice displayed resilience, as 57.1% of males and 60% of females were resilient. A Trend LogRank for escapes (censored for no escapes) indicated a faster escape performance in resilient mice compared to the helpless group (Fig. 1D; X^2^: 65.53, df = 1, p < 0.0001).

**Figure 1.**
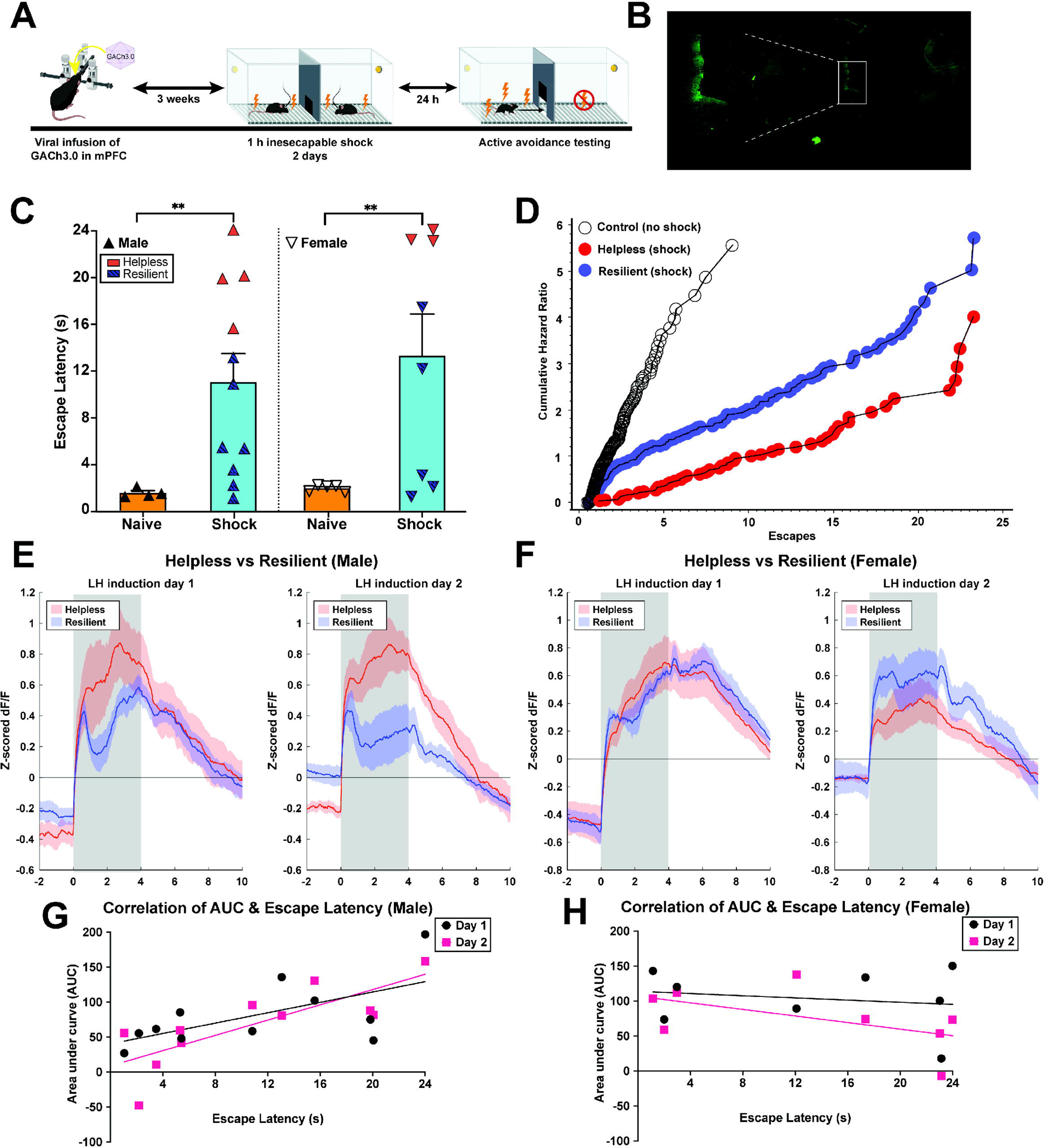
ACh levels in the mPFC are predictive of escape behavior in the learned helplessness paradigm. **A.** Experimental design. Virus containing a construct for the fluorescent ACh sensor GACh3.0 was infused into the mPFC of male and female WT mice. Mice were allowed to recover for 3 weeks prior to beginning learned helplessness training, which consisted of two successive days of 1 h induction sessions of 120 inescapable shocks each. On the third day, mice underwent active avoidance training during which shocks and the possibility to escape were presented simultaneously. **B.** Image of mPFC and a higher magnification inset with fiber tract and GACh3.0 infection (green fluorescence surrounding fiber tract). **C.** Exposure to shocks during induction sessions resulted in increased escape latencies in male (left) and female (right) mice. Using a *k-means* clustering algorithm, mice were divided into helpless and resilient groups based on both their escape latency and number of trials with a failure to escape during active avoidance testing. **D.** A Trend LogRank for escapes (censored for no escapes) indicated a significant difference in escape performance between helpless and resilient mice, in which resilient mice achieved full escape efficacy sooner than helpless mice. **E.** Helpless males displayed increased ACh signaling in response to 4-s shock (grayed area) during induction trials, (**F**) whereas female helpless and resilient mice displayed similar ACh signaling. Data shown are the average z-scored ΔF/F per group, per day. **G.** There was a significant correlation between average AUC of ACh signal in response to shock during induction trials and escape latency during subsequent active avoidance testing for individual male mice on both days 1 and 2, but not for female mice (**H**).

A robust ACh increase was observed in the mPFC in response to onset of the 4-s shock, remained elevated until termination, and did not return to baseline until ∼8-10 s post-shock termination (Supplementary Fig. 1C). Furthermore, helpless and resilient mice showed different patterns of ACh response during learned helplessness training, with helpless males showing larger ACh responses to shock (Fig 1. E,F). Furthermore, although rates of helplessness did not differ between male and female mice, the area under the curve (AUC) of the cholinergic signal measured during induction sessions was positively corelated with escape latency only in males (Fig. 1G, H; Day 1: r = 0.61, p < 0.05; Day 2: r = 0.79, p < 0.01), suggesting that there is a sex difference in the contribution of mPFC ACh to escape responses following inescapable shock.

### Pharmacologically increasing ACh during LH induction increases escape deficits in later active avoidance testing

To determine whether the large difference in mPFC ACh levels between helpless and resilient mice on day 2 of induction contributed to the active avoidance deficit, we administered physostigmine i.p. 30 min prior to the second induction session (Fig. 2A). Because repeated physostigmine administration can increase AChE levels (*25*), we chose to administer the drug only on Day 2 of induction, given the robust correlation between ACh signaling on Day 2 and escape latency observed in our fiber photometry experiment. Previous studies have shown that i.p. physostigmine injection decreases AChE activity in the PFC, along with other brain regions (*13*). No sex differences were detected, so data were pooled for male and female mice.

**Figure 2.**
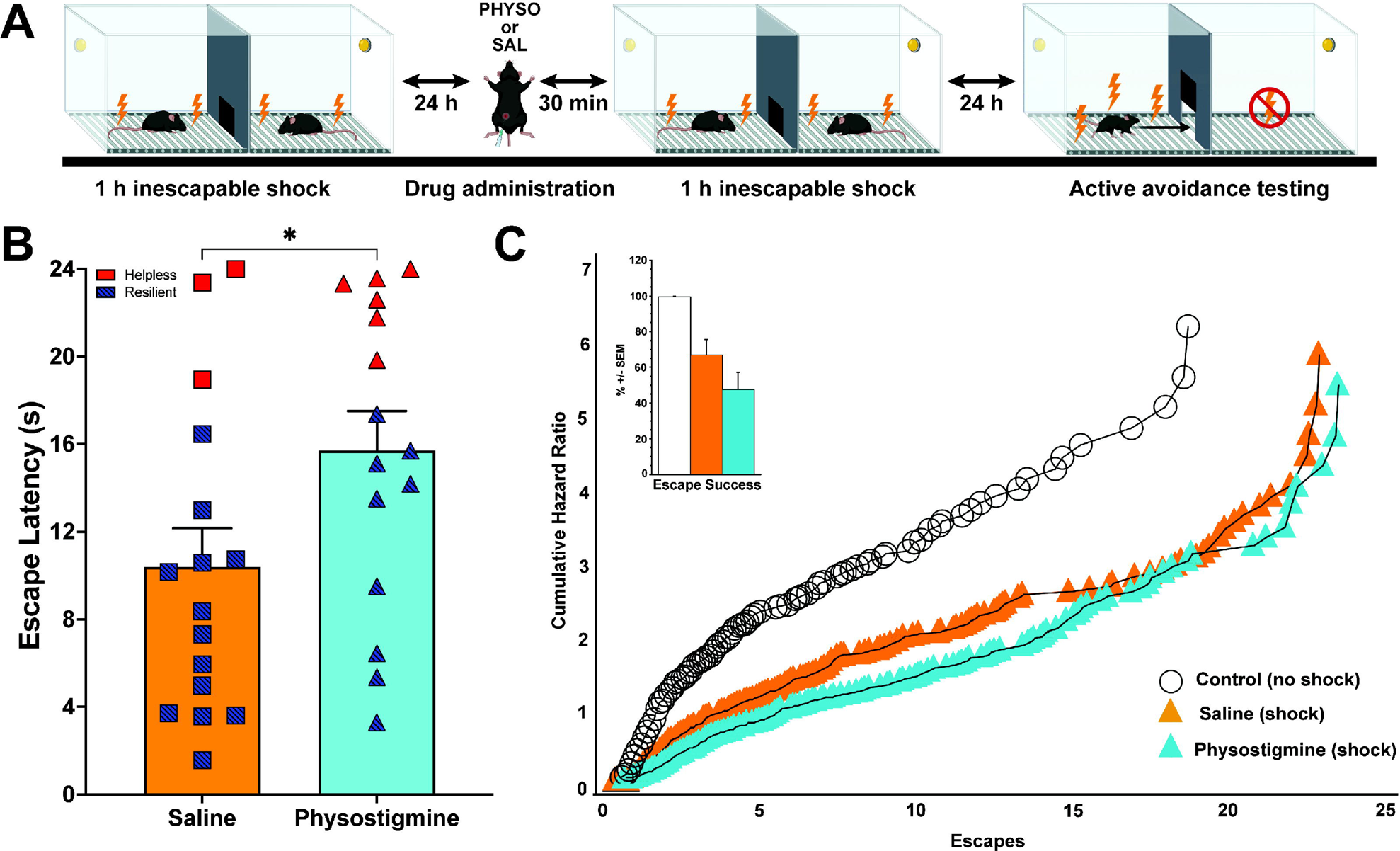
Pharmacologically prolonging ACh signaling during LH induction increases escape deficits in later active avoidance testing. **A.** Experimental design. Mice received injections of physostigmine (PHYSO) or saline (SAL) 30 min prior to their second induction session. **B.** Physostigmine administration increased escape latencies. **C.** Relative to saline, physostigmine decreased escapes in the active avoidance test (**insert**). Furthermore, a Trend LogRank for escapes (censored for no escapes) indicated a significant difference in escape performance between mice that received physostigmine and those that received saline, such that saline mice achieved full escape efficacy sooner than those treated with physostigmine.

Physostigmine administration during induction on Day 2 increased escape latencies in subsequent active avoidance trials (Fig. 2B; t (29) = 2.10, p < 0.05). There were no differences in helpless-to-resilient ratios between saline and physostigmine groups (3 of 16 saline helpless mice and 6 of 16 physostigmine helpless (*X^2^* (1, N = 32) = 1.39, p = 0.24). Relative to saline, physostigmine decreased the number of successful escapes in the active avoidance test (Fig. 2C insert; Mann-Whitney *U* = −5.16, *n*_1_ = 16 *n*_2_ = 15, *P* < 0.001). Furthermore, a Trend LogRank for escapes (censored for no escapes) indicated a significant difference in escape performance between mice that received physostigmine and those that received saline, in which saline animals achieved full escape efficacy sooner than those treated with physostigmine (Fig. 2C; *x*^2^ :135.9, df = 1, p < 0.0001).

### Manipulation of ACh signaling in mPFC during LH training using DREADD constructs

ACh signaling is increased in the mPFC of helpless male mice and pharmacological manipulation of ACh levels in both sexes during induction trials resulted in escape deficits in active avoidance testing. We therefore determined whether selectively manipulating ACh inputs to the mPFC during induction trials was sufficient to influence escape performance in the active avoidance test. We infused retrograde AAVs carrying Cre-dependent DREADD constructs into the mPFC of ChAT-Cre mice to either inhibit (Gi) or excite (Gq) cholinergic neurons projecting to the mPFC of mice during learned helplessness induction (Fig. 3A).

**Figure 3.**
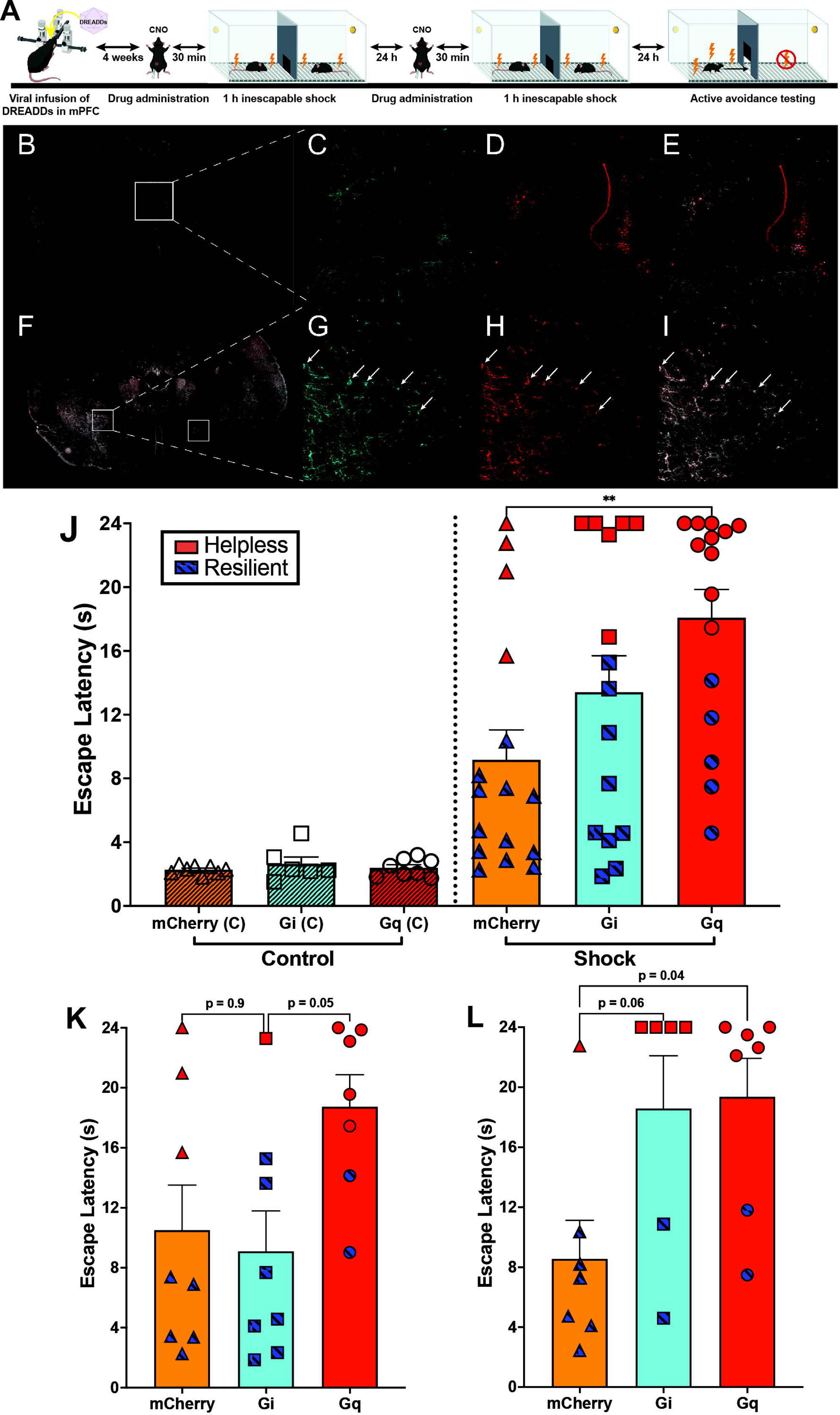
Effects of DREADD-mediated manipulation of cholinergic terminals in the mPFC. **A.** Experimental design. Retrograde, Cre-dependent viral constructs expressing Gq- or Gi-DREADDs or mCherry control were infused into the mPFC of ChAT-Cre mice. 4 weeks later mice underwent learned helplessness induction and active avoidance testing. Mice were administered CNO 30 minutes prior to each induction session. **B.** Brain section containing mPFC, the target for viral infusion. **C, D, E.** Higher magnification image of mPFC showing ChAT in cyan (**C**), DREADD expression in mCherry (**D**), and both signals merged (**E**). **F.** Brain section containing nucleus basilis cell bodies infected with DREADD-containing retrograde virus injected into mPFC. **G, H, I.** Higher magnification image of nucleus basilis showing cell bodies (arrows) infected with DREADD-containing virus, with ChAT in cyan (**G**), DREADD in mCherry (**H**), and both signals merged (**I**). **J.** Shock exposure during induction increased escape latency in all groups relative to controls. Gq-mediated excitation of ChAT-expressing terminals in the mPFC during induction increased escape latency in active avoidance testing. **K.** In male mice, Gq-mediated excitation increased escape latencies relative to Gi-mediated inhibition, but not relative to controls. Gi-mediated inhibition also resulted in more resilient mice than those that received Gq-mediated excitation. **L.** In contrast, relative to controls, female mice showed increased escape latencies in active avoidance testing following both Gq-mediated excitation and Gi-mediated inhibition during induction trials.

Escape latencies for mCherry control mice exposed to shock showed significantly increased escape latencies compared to non-shocked mCherry control mice, as expected (Fig. 3J; mean diff = 7.46 s, t (20) = 2.60, p < 0.05). Approximately 73% of mCherry control mice in the shock group were resilient. These data demonstrate that surgery and viral infusion alone did not alter helplessness behavior compared to unmanipulated mice.

Gq-mediated excitation of cholinergic neurons projecting to the mPFC was sufficient to increase escape latencies in an active avoidance test (Fig. 3J; F (5, 62) = 11.51, p<0.001) and increase helplessness among shock mice (*x^2^* (1, N=31)=5.43, p<0.05). No differences were observed between control groups. Although helplessness-to-resilient ratios of Gi- or Gq-expressing mice did not differ from mCherry controls (*x^2^* (1, N=16)=1.33, p=0.25 and *x^2^*(1, N=15)=1.73, p=0.19, respectively), Gi-mediated inhibition of cholinergic neurons in males increased resilience relative to Gq-mediated excitation (*x^2^* (1, N=15)=5.40, p<0.05). ANOVA indicated a main effect of DREADD (Fig. 3J; F (2, 20) = 3.59, p<0.05) with *post hoc* comparisons revealing that Gq-mediated excitation increased escape latencies in active avoidance testing relative to Gi-mediated inhibition (p=0.05).

In female mice there was a significant main effect of DREADD type (Fig 3L; F (2, 17) = 4.57, p<0.05) with *post hoc* comparisons indicating that Gq-mediated excitation of mPFC cholinergic projections significantly increased escape latencies compared to control female mice. Paradoxically, there was also a strong trend for an increase in escape latencies due to Gi-mediated inhibition of mPFC cholinergic inputs (p=0.06) relative to mCherry controls. Both Gi-DREADD-mediated inhibition and Gq-DREADD-mediated excitation resulted in increased helplessness relative to mCherry controls (*x^2^* (1, N=15)=5.40, p<0.05 and *x^2^* (1, N=16)=4.27, p<0.05, respectively) but did not differ from each other (*x^2^* (1, N=15)=0.13, p=0.36). Along with data from fiber photometry experiments, this indicates that mPFC cholinergic signaling regulates escape behavior differently in male and female mice, with female mice showing optimal escape behavior under control conditions, whereas escape behavior is disrupted as a result of either increasing or decreasing mPFC ACh levels during training.

## Discussion

The data shown here demonstrate that ACh signaling in the mPFC of mice is critical in setting the balance between susceptibility and resilience to helplessness following exposure to inescapable stressors. ACh dynamics in the mPFC during presentation of identical inescapable shocks predicted failure to escape during an active avoidance task. Concordantly, increasing activity of ACh inputs to the mPFC during induction trials induced escape deficits in later active avoidance testing and increased the proportion of helpless-to-resilient mice. In addition, we identified sex differences in how ACh dynamics during inescapable shock predict escape behavior, and how chemogenetic manipulation of ACh activity drives helplessness. In male, but not female, mice there was a positive correlation between ACh activity and escape latency in active avoidance trials. However, both pharmacological and chemogenetic increases in ACh activity were sufficient to increase escape latency and helplessness in mice of both sexes. In contrast, inhibiting ACh activity in the mPFC reduced escape deficits and conferred resilience in male mice, but increased escape deficits and helplessness in female mice. These data suggest that resilience to inescapable stress can be conferred in male mice by decreasing mPFC ACh levels, but that ACh levels are optimal in female mice at baseline, and either increasing or decreasing levels results in greater susceptibility to stress.

Using fiber photometry with the GRAB ACh 3.0 sensor we observed an increase in ACh signaling in the mPFC in response to shock. This is consistent with microdialysis studies showing increased ACh in the PFC during exposure to adverse stimuli including restraint and inescapable shock (*26*). Compared to the timescale of microdialysis (many minutes), fiber photometry provides data on a subsecond scale, allowing us to observe how each 4-s shock affects ACh fluctuations. ACh signaling increased immediately upon shock presentation, with sustained signaling throughout the shock, followed by a slow decay 4-6 s post-shock termination to return to baseline. These dynamics are similar to those observed in the hippocampus, with some important differences. Following a 1-s shock, ACh levels in the ventral hippocampus not only increased during shock but continued to increase for ∼1 s post shock termination before taking a further ∼15 s to return to baseline, an even slower decay than observed in the mPFC. In the dorsal hippocampus, ACh signaling increased during the shock but began to decrease immediately upon shock termination, returning to baseline ∼4-5 s post shock (*27*). Signaling in the basolateral amygdala also follows a different pattern, such that 1-s shocks elicit an immediate increase in ACh signaling that returns to baseline within ∼1 s post shock termination (*20*). It is important to note that AChE activity differs in these regions (*28*), and this is likely to alter the dynamics of ACh clearance across these brain areas.

Several studies have shown that manipulating ACh transmission significantly alters stress-related behaviors. Decreasing AChE activity in the hippocampus increases immobility in the tail-suspension test (TST; a form of passive coping) and promotes susceptibility to social stress (*9*), whereas knockdown of β2-containing and α7 nicotinic ACh receptors (nAChR) in the amygdala increases resilience to social defeat stress and increased active coping in TST and forced swim test (FST) (*12*); similarly administration of the α7-containing nAChR agonist methyllycaconitine decreases immobility in the TST and the FST (*29*). Consistent with the current study showing that decreasing mPFC ACh signaling results in resilience to inescapable stress, the antidepressant-like effects of the muscarinic antagonist scopolamine are dependent upon M1-type muscarinic ACh receptors in the mPFC (*30–32*). The current study is therefore consistent with the observation that depressed individuals exhibit hyperactivity of the cholinergic system and that this may contribute to greater stress-susceptibility.

Along with its role in behavioral responses to stress, ACh in the mPFC is also recruited as a result of effortful processing, both in tests of attention and other cognitive tasks. Depletion of ACh in the PFC of primates results in deficient spatial working memory, while activating α7 nAChRs improves primate working memory (*33*). Cholinergic signaling in the mPFC drives attention toward salient sensory information while increases in the region also enhance salient cue detection (*17, 34*). Cholinergic signaling follows an inverted U-shaped curve however, in which both too much or too little ACh is detrimental to cognitive processes and behavioral outcomes (*35*). For instance, use of either a selective muscarinic type-1 receptor agonist or a positive allosteric modulator can improve working memory at low doses but impair it at higher doses (*36*). This inverted-U shaped function suggests that an optimal range of ACh activity is beneficial, while moving significantly higher or lower than this can impair cognitive function. This is similar to the impairment of active avoidance response we observed here in female mice following either activation or inhibition of mPFC ACh inputs during exposure to inescapable stress. Impairment of cognitive function dependent on activity of the PFC occurs following exposure to uncontrollable stress (*37, 38*) and could therefore contribute to the helplessness phenotype observed here.

At baseline, both male and female mice appear to be at an optimal mPFC ACh level to avoid helplessness. However, in both male and female mice, increasing ACh above this optimum may lead to overly strong encoding of the shock-environment association during inescapable shock, thereby impairing behavioral flexibility during exposure to escapable shock in the active avoidance test. Prolonged elevated ACh signaling in males and females could increase attention to the environmental cues perceived during presentation of inescapable shocks, allowing overly potent encoding that leads to inability to learn a new response when the shocks become escapable. Scopolamine exerts its antidepressant properties via M1-type muscarinic signaling within the PFC (*30–32*). M1 signaling onto mPFC pyramidal neurons may therefore be important for optimal tuning of these neurons, as has been shown for α7 nAChRs (*39*) and dopamine D1 signaling (*40*).

Decreased mPFC ACh signaling resulted in opposing effects in males and females, with males experiencing increased resilience to the inescapable shocks and females showing greater susceptibility to helplessness. This divergence should perhaps not be surprising given that stress produces sexually dimorphic effects in the mPFC. Although acute, uncontrollable stress is known to weaken synaptic function in the PFC (*41*), stress has been shown to induce dendritic neuronal atrophy and decreasing apical dendritic branch length in the mPFC of male rats, while either having no effect on or, in some cases, increasing dendritic length in female rats (*42–44*). Interestingly, baseline PFC paired-pulse facilitation – a form of short-term synaptic potentiation – may be greater in female mice than in males (*45*). This could indicate more robust neuronal responding in response to repeated stimuli in the mPFC of female mice compared to male mice. ACh signaling could contribute to this form of potentiation, since nAChRs are important regulators of synaptic plasticity in the mPFC (*46*). It is important to note, however, that despite these physiological differences, similar rates of resilience were observed in both sexes, suggesting that similar behavioral outcomes can emerge from divergent physiological mechanisms. This is in line with studies showing that compounds with antidepressant properties can have similar behavioral effects in male and female mice but little overlap in their neurobiological effects at a circuit level (*47*).

The studies shown here suggest that ***increases*** in mPFC ACh levels during the induction phase of LH may increase attention to environmental cues and contingencies during exposure to stressful, inescapable shocks, leading to greater recruitment of memory circuits that strengthen encoding of these negative stimuli, resulting in maladaptive passive coping during the active avoidance trial. For female mice, a ***decrease*** in mPFC ACh signaling also results in increased helplessness, and this may reflect a different type of learning deficit that requires ACh recruitment for flexible behavior. Taken together, the current findings provide strong support for a role of ACh in a negative encoding bias, suggesting that homeostatic levels of ACh signaling are necessary for appropriate learning and affect, but overly high or prolonged elevation of cholinergic signaling promote encoding of stressful events, and could lead to rumination in human subjects, a core symptom of depression (*10*).

## Acknowledgements

These studies were supported by grants MH077681, MH105824 and DA033945 from the National Institutes of Health. This work was funded in part by the State of Connecticut, Department of Mental Health and Addiction Services, but this publication does not express the views of the Department of Mental Health and Addiction Services or the State of Connecticut.

