## Supplemental Figure 1 for "Acetylcholine signaling in the medial prefrontal cortex mediates the ability to learn an active avoidance response following learned helplessness training"

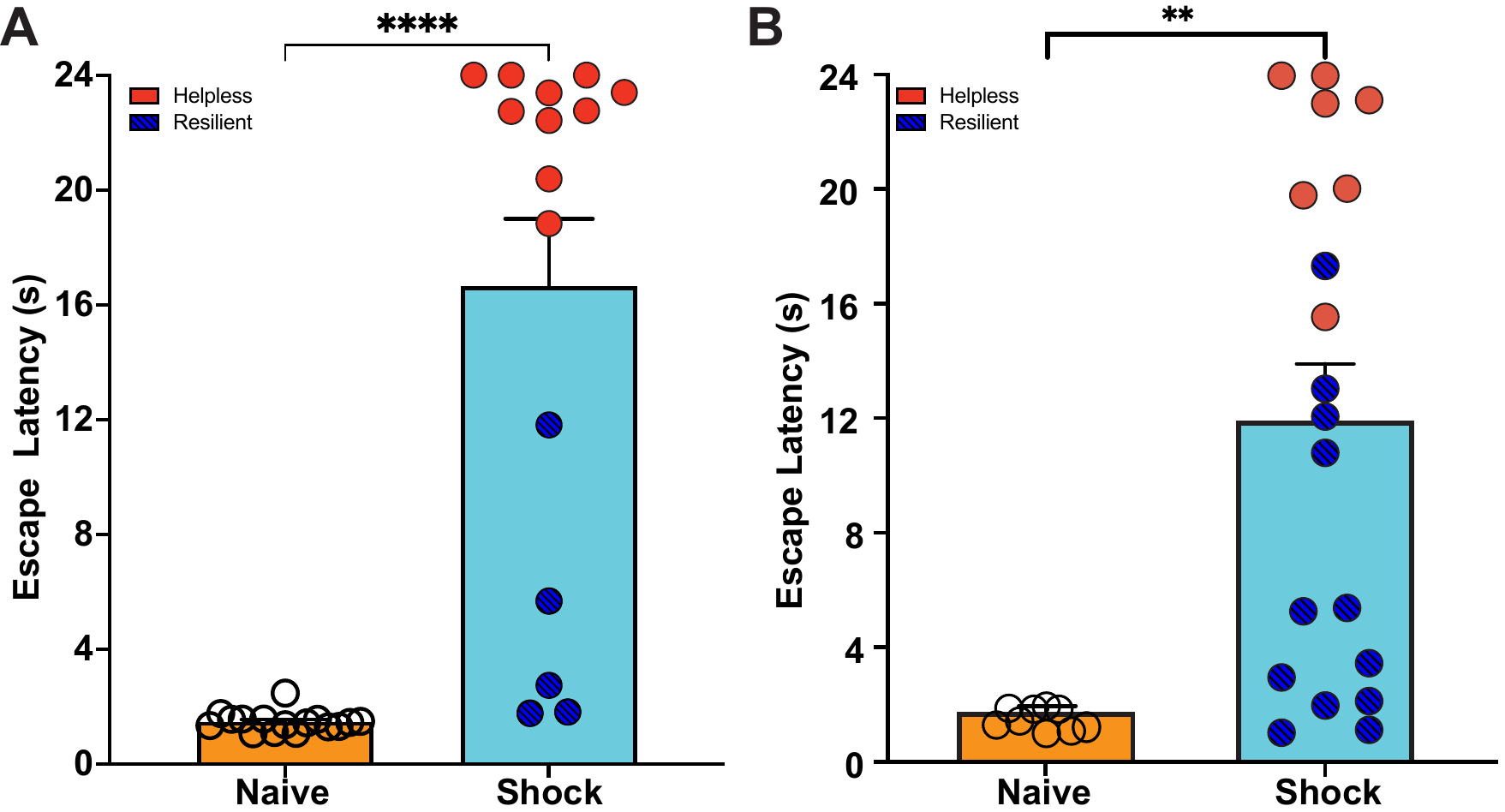

**Supplemental Figure 1**. **A.** Wildtype mice that received shocks during induction sessions in the learned helplessness paradigm showed significantly elevated escape latencies in active avoidance testing relative to control animals that did not receive shocks during induction sessions. **B.** In the fiber photometry experiment, mice in the shock group displayed elevated escape latencies relative to mice that also underwent surgery but were naïve to shock.
